# HDAC11 Suppresses the Thermogenic Program of Adipose Tissue via BRD2

**DOI:** 10.1101/292011

**Authors:** Rushita A. Bagchi, Bradley S. Ferguson, Matthew S. Stratton, Tianjing Hu, Maria A. Cavasin, Lei Sun, Ying-Hsi Lin, Dianxin Liu, Pilar Londono, Kunhua Song, Maria F. Pino, Lauren M. Sparks, Steven R. Smith, Philipp E. Scherer, Sheila Collins, Edward Seto, Timothy A. McKinsey

**Affiliations:** Department of Medicine, Division of Cardiology, University of Colorado Anschutz Medical Campus, Aurora, CO 80045, USA; Consortium for Fibrosis Research & Translation, University of Colorado Anschutz Medical Campus, Aurora, CO 80045, USA; George Washington University Cancer Center, Washington, DC 20037, USA; Integrative Metabolism Program, Sanford Burnham Presbys Medical Discovery Institute at Lake Nona, Orlando, FL 32827, USA; Translational Research Institute for Metabolism and Diabetes, Florida Hospital, Orlando, FL 32827, USA; Touchstone Diabetes Center, Department of Internal Medicine, University of Texas Southwestern Medical Center, Dallas, TX 75390, USA

## Abstract

Little is known about the biological function of histone deacetylase 11 (HDAC11), which is the lone class IV HDAC. Here, we demonstrate that deletion of HDAC11 in mice stimulates brown adipose tissue (BAT) formation and beiging of white adipose tissue (WAT). Consequently, HDAC11-deficient mice exhibit dramatically enhanced thermogenic potential and, in response to high fat feeding, attenuated obesity, insulin resistance, and hepatic steatosis. *Ex vivo* and cell-based assays revealed that HDAC11 catalytic activity suppresses the BAT transcriptional program, in both the basal state and in response to β-adrenergic receptor signaling, through a mechanism that is dependent on physical association with BRD2, a bromodomain and extraterminal (BET) acetyl-histone binding protein. These findings define a novel epigenetic pathway for the regulation of energy homeostasis, and suggest potential for HDAC11-selective inhibitors for the treatment of obesity and diabetes.

## Introduction

It is estimated that greater than 1/3 of the population of the United States is obese (1). Obesity is coupled to the development of many chronic diseases, including type 2 diabetes (T2D), which is projected to afflict over half a billion adults worldwide by 2040 (2). Obesity is a consequence of a disparity between energy intake and expenditure (3), and decreasing obesity requires a reduction in energy intake and/or an increase in energy expenditure. Aside from diet and exercise, therapeutic interventions for obesity include bariatric surgery and the use of drugs that decrease energy intake (4).

There is intense interest in developing alternative pharmacotherapy for obesity based on increasing energy expenditure via brown adipose tissue (BAT) (5). In contrast to white adipose tissue (WAT), which functions mainly to store energy in the form of triglycerides in unilocular white adipocytes, brown adipocytes within BAT harbor small, multilocular lipid droplets and an abundance of mitochondria, which produce heat through non-shivering thermogenesis (6). Heat production by BAT is governed by uncoupling protein-1 (UCP1), which resides in the inner mitochondrial membrane in brown adipocytes and functions as a long-chain fatty acid/H+ symporter to catalyze mitochondrial proton leak and thereby uncoupling electron transport from ATP synthesis (7-9). BAT is highly metabolically active, and has been shown to contribute to energy expenditure in humans (10). Additional studies in humans have revealed that body mass index and percent body fat negatively correlate with BAT abundance (11), and a polymorphism in the gene encoding UCP1 is associated with fat gain and obesity (12). Together, these findings validate the potential of BAT-targeted therapies for the treatment of obesity.

Pharmacological approaches to promote BAT formation and function have included the use of β_3_-adrenergic receptor (β_3_-AR) agonists. β_3_-AR stimulation directly enhances lipolysis and energy expenditure, and also triggers downstream signaling events that lead to induction of a thermogenic gene program, which includes the gene encoding UCP1 (13). Furthermore, β_3_-AR stimulation of WAT can trigger the emergence of UCP1-expressing cells, termed beige or brite fat (14;15), with morphological and functional characteristics of brown adipocytes (5;16;17). However, while β_3_-AR agonists were shown to acutely increase energy expenditure and insulin sensitivity in humans, they failed to promote weight loss upon chronic administration (18-21). Other approaches have targeted transcriptional regulators, primarily nuclear hormone receptors, which stimulate thermogenic gene expression. For example, peroxisome proliferator-activated receptors (PPARs) and thyroid hormone receptors function downstream of the β_3_-AR to promote BAT gene expression and beiging of WAT, and PPAR agonists and thyromimetics have been shown to be efficacious in animal models of obesity (22-30). However, it is unclear whether pharmacological activation of these transcription factor pathways will be sufficiently tolerated in humans to provide a viable avenue for treatment of obesity (31).

Here, using multiple *in vivo*, *ex vivo* and cell-based approaches, we demonstrate that histone deacetylase 11 (HDAC11) functions as a repressor of the thermogenic gene program in BAT, and prevents beiging of WAT. Compared with wildtype controls, mice lacking HDAC11 are lean and harbor excess BAT. Accordingly, HDAC11 deficiency leads to enhanced cold-induced thermogenesis, reduced weight gain and lipid accumulation in response to high fat feeding, and improved glucose tolerance. The global metabolic effects of HDAC11 deletion correlate with enhanced UCP1 expression in BAT, a profound increase in beiging of WAT, and augmented thermogenic gene expression in response to β_3_-AR signaling. Using cell-based models, we provide evidence for cell-autonomous roles for HDAC11 as a repressor of brown adipocyte differentiation and thermogenic gene expression, functions that are dependent on association of HDAC11 with the bromodomain and extraterminal (BET) family member BRD2. These data demonstrate a critical and previously unrecognized role for HDAC11 as an epigenetic regulator of whole-body metabolism. Furthermore, since HDAC11-deficient mice are healthy (32), and HDAC11 has a unique catalytic domain compared to other HDAC isoforms (33), the findings suggest the intriguing possibility that selective HDAC11 inhibitors could be developed to increase energy expenditure for the treatment of obesity and metabolic disease.

## Results

### HDAC11 deficiency increases BAT abundance and function, and triggers beiging of WAT

To further define the biological functions of HDAC11, we began investigating mice in which the *Hdac11* gene was deleted globally (KO). KO mice were indistinguishable from wildtype controls (WT) with regard to overt morphological characteristics (Figure 1A). However, necropsy revealed increased interscapular brown adipose tissue (iBAT) mass accompanied by a significant reduction in inguinal white adipose tissue (ingWAT) and epididymal WAT (eWAT) mass in KO mice compared to WT littermates (Figure 1, B-D). Consistent with a role for HDAC11 in the differential control of distinct fat depots, *Hdac11* mRNA was more highly expressed in WAT compared to BAT in mouse and human samples (Figure 1, E and F). These findings suggest a previously unrecognized role for HDAC11 in the control of adipose tissue.

**Figure 1.**
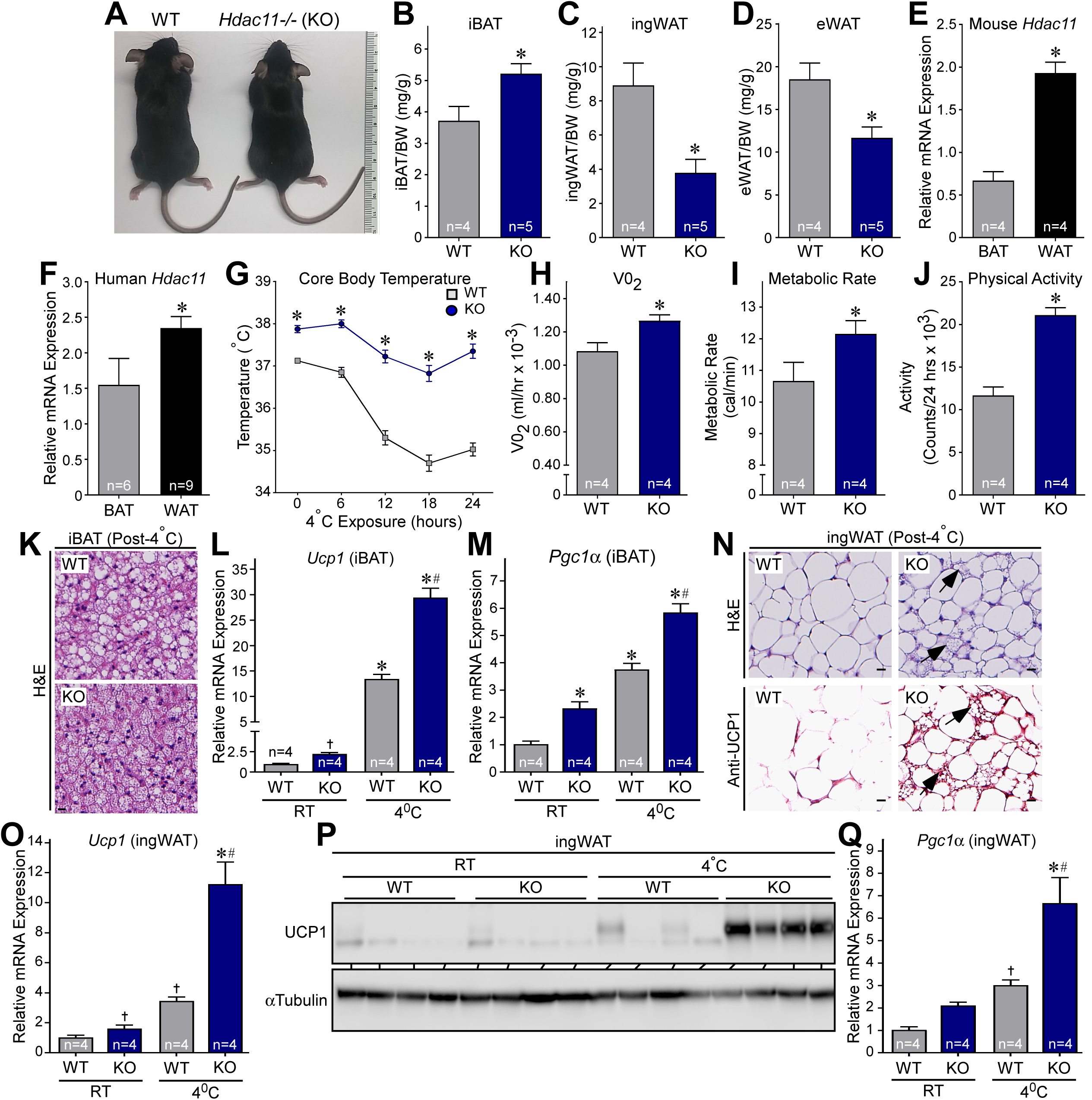
HDAC11 deficiency increases thermogenic potential in mice. (**A**) Gross morphology of 10-week-old male wildtype (WT) and *Hdac11-/-* (KO) littermates fed standard rodent chow. (**B-D**) Interscapular brown adipose tissue (iBAT), inguinal white adipose tissue (ingWAT) and epididymal WAT (eWAT) from 10-week-old male mice were dissected and weighed. (**E** and **F**) Quantitative PCR analysis of *Hdac11* mRNA expression in mouse and human BAT and WAT. (**G**) WT and KO mice were fed standard rodent chow, exposed to 4°C for 24 hours, and core body temperature was recorded at the indicated time points using a rectal thermal probe; n = 4 mice/group. (**H-I**) Oxygen consumption (VO_2_) and metabolic rate (MR) were measured using a CLAMS system for 24 hours. (**J**) Activity was calculated using laser beam breaks per 24 hours. (**K**) H&E staining of iBAT post-24 hours of 4°C exposure. Scale bar:10 μm. (**L-M**) Quantitative PCR analysis of *Ucp1* and *Pgc1*α mRNA expression in iBAT from mice housed at room temperature (RT) or exposed to 4°C for 24 hours. (**N**) H&E staining and UCP1 immunohistochemistry of ingWAT 24 hours post-4°C exposure. Arrows indicate evidence of beiging. (**O**) Quantitative PCR analysis of *Ucp1* mRNA expression in ingWAT. (**P**) Immunoblot analysis of UCP1 protein expression in ingWAT. (**Q**) Quantitative PCR analysis of *Pgc1*α mRNA expression in ingWAT. Values for all graphs represent means +SEM; \**P*<0.05 vs. WT; ^#^*P*<0.05 vs. WT 4°C using ANOVA with Tukey’s multiple comparisons test. Student’s t-test was employed for two groups, and ^†^*P*<0.05 vs WT RT using unpaired t-test following significant ANOVA.

Given the function of BAT in mediating non-shivering thermogenesis, mice were subsequently placed in metabolic cages and core body temperature and metabolic parameters were measured at ambient temperature (22 °C) and during a 24-hour cold challenge at 4 °C. Compared to WT controls, KO mice had elevated body temperature prior to and during the cold challenge, which correlated with increased oxygen consumption, metabolic rate and physical activity (Figure 1, G-J). Consistent with this, histological analysis following cold exposure revealed smaller lipid droplets in iBAT from KO mice compared to WT controls, which is a characteristic of accelerated triglyceride liberation (Figure 1K). Furthermore, expression of BAT-selective thermogenic genes, including *Ucp1* and *Pgc1α,* was also increased in KO iBAT compared to WT controls (Figure 1, L and M). Strikingly, subcutaneous ingWAT from KO mice subjected to cold challenge exhibited marked signs of beiging, including increased abundance of multi-lobular adipocytes, pronounced induction of UCP1 protein expression, and increased levels of *Ucp1* and *Pgc1α* mRNA expression (Figure 1, N-Q); qPCR analysis confirmed the absence of *Hdac11* mRNA transcripts in KO adipose tissue (Supplemental Figure 1). Together, these data suggest that BAT function is ameliorated by the absence of HDAC11, and that deletion of HDAC11 promotes beiging of WAT.

Induction of thermogenic gene expression upon cold challenge is governed by β_3_-AR signaling. To begin to address whether HDAC11 functions in an adipose tissue-autonomous manner to control thermogenesis, acute *ex vivo* studies were performed with iBAT explants from WT and KO mice (Figure 2A). Induction of *Ucp1* and *Pgc1α* mRNA expression by the β_3_-AR-selective agonist, CL-316,243, was significantly potentiated in KO iBAT compared to WT controls (Figure 2, B and C). Although expression of β_3_-AR mRNA expression was also elevated in KO iBAT following agonist treatment, baseline expression of the receptor was equivalent in WT and KO iBAT (Figure 2D). These data suggest that HDAC11 functions, at least in part, within adipose tissue to repress thermogenic β_3_-AR target gene expression, rather than by generally increasing adrenergic signaling.

**Figure 2.**
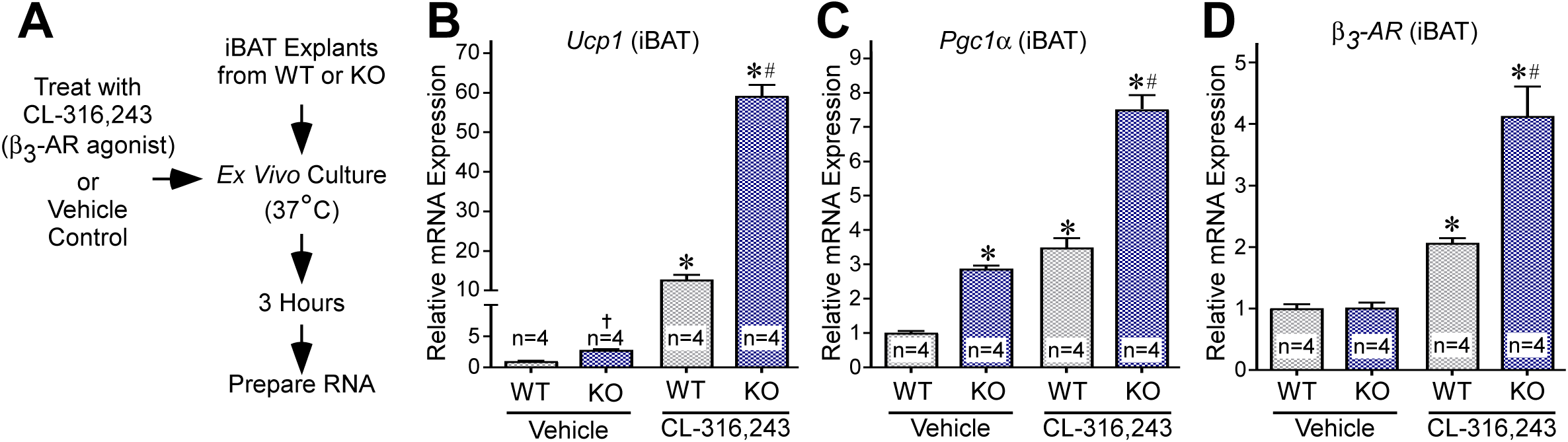
HDAC11 deficiency increases β_3_-AR-stimulated gene expression in BAT *ex vivo*. (**A**) Schematic depiction of the iBAT *ex vivo* signaling assay. (**B-D**) Quantitative PCR analysis of *Ucp1*, *Pgc1*α and β_3_-*AR* mRNA expression in iBAT treated *ex vivo* with CL-316,243 or vehicle control (DMSO; 0.1% final concentration) for 3 hours. Values for all graphs represent means ^+^SEM; \**P*<0.05 vs. WT Vehicle; ^#^*P*<0.05 vs. WT CL-316,243 using ANOVA with Tukey’s multiple comparisons test. ^†^*P*<0.05 vs WT Vehicle using unpaired t-test following significant ANOVA.

### HDAC11 deficiency protects against deleterious effects of high fat feeding

Activation of BAT and beiging of WAT can attenuate obesity and metabolic dysfunction. To initially assess the impact of deleting HDAC11 on metabolic homeostasis, WT and KO mice were individually housed in metabolic cages, fed a high fat, high sucrose diet (HFD), and monitored continuously over an eight day period during ad libitum feeding (Figure 3A). Remarkably, despite equivalent food intake between WT and KO mice, weight gain was reduced by ~50% in KO mice (Figure 3, B and C). Relative to WT controls, KO mice exhibited increased oxygen consumption as well as higher metabolic rate and total energy expenditure, suggesting enhanced fatty acid oxidation in KO adipose tissue (Figure 3, D-F). Glucose tolerance was also enhanced in KO mice compared to WT controls at baseline and following high fat feeding (Figure 3, G and H). Quantitative magnetic resonance imaging (QMRI) revealed that the overall improvement in systemic metabolism in KO mice correlated with reduced fat mass post-HFD, and elevated lean mass before and after HFD compared to WT controls (Figure 3, I and J).

**Figure 3.**
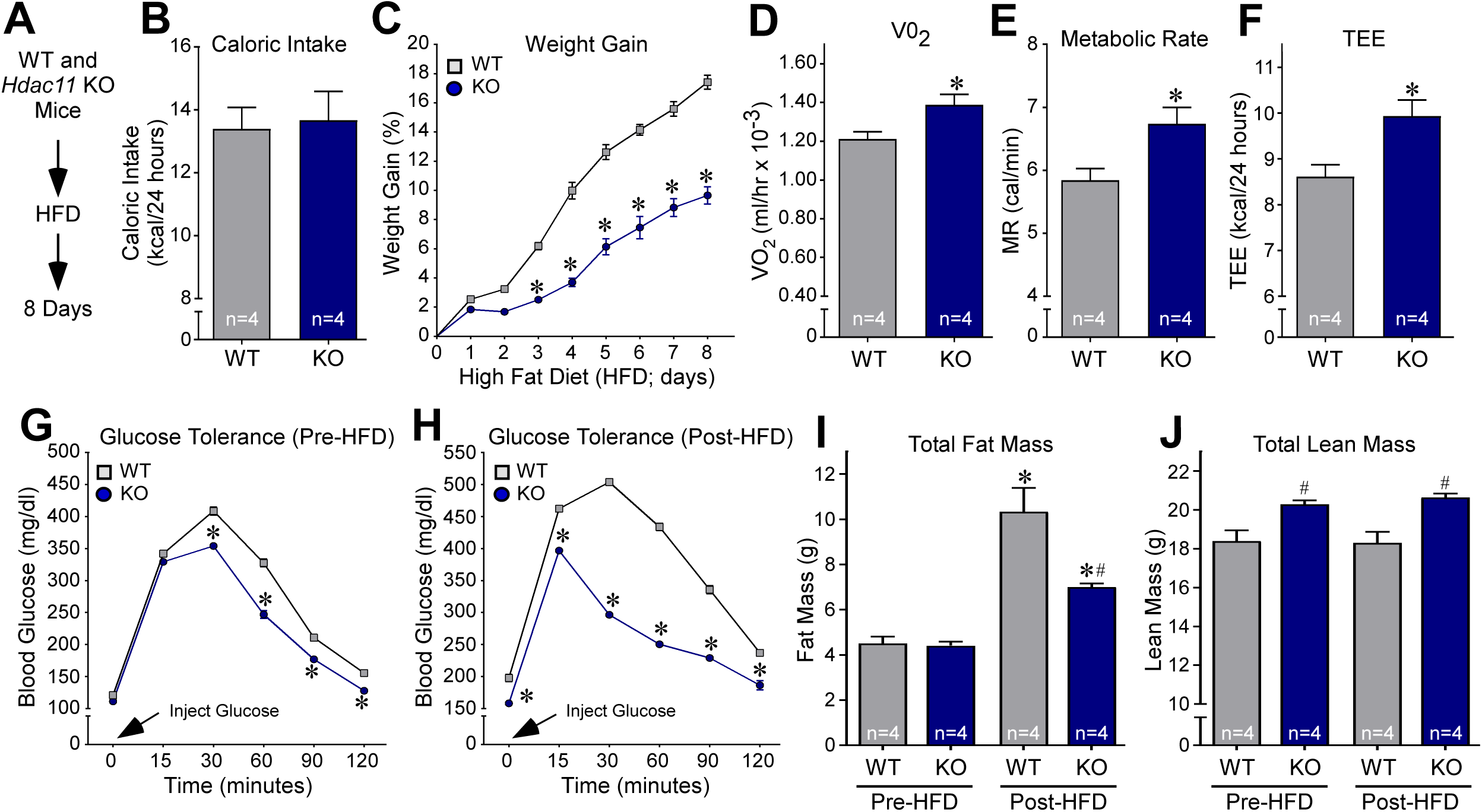
HDAC11 deficiency increases metabolism and glucose tolerance in response to acute high fat feeding. (**A**) Schematic representation of the high fat diet (HFD) experiment. (**B**) Daily food consumption was determined using an automated food hopper attached to CLAMS chambers. (**C**) Weight gain (%) was calculated each day for 8 days. n = 4 animals/group; \**P*<0.05 by ANOVA. (**D-F**) Oxygen consumption (VO_2_), metabolic rate (MR) and total energy expenditure (TEE) were measured using the CLAMS system. \**P*<0.05 by Student’s t-test. (**G**) Glucose tolerance was assessed in mice prior to high fat feeding; n = 4/group. (**H**) Glucose tolerance was assessed in mice following 8 days of high fat feeding; n = 4/group. (**I** and **J**) Body composition (fat and lean mass) was determined using QMRI. Values for all graphs represent means +SEM; (**C-H**) *P<0.05 vs. WT; (**I** and **J**) \**P*<0.05 vs. WT Pre-HFD and ^#^*P*<0.05 vs. WT Post-HFD. Two-way ANOVA with Sidak’s or Tukey’s multiple comparisons test for four groups was employed for (**C**, **G**, **H**) and (**I** and **J**) respectively; Student’s t-test was used for (**B**, **D-F**).

To determine whether HDAC11 deficiency confers chronic protection from deleterious effects of high fat feeding, WT and KO mice were fed a HFD for 12 months (Figure 4A). Consistent with findings from the acute study, weight gain was blunted, and glucose tolerance enhanced in KO mice compared to WT controls (Figure 4, B and C). These improvements in KO mice were associated with reduced circulating levels of insulin and leptin, and a concomitant increase in circulating adiponectin, a fat-derived hormone that is negatively correlated with metabolic dysfunction (Figure 4, D-F). QMRI demonstrated significantly lower fat mass in KO mice fed HFD compared to WT controls, with no change in lean mass between the groups (Figure 4, G and H). In line with the lower fat mass in KO mice, histological analyses revealed small lipid droplets in iBAT, reduced size of triglyceride storing adipocytes in eWAT, and attenuated hepatic steatosis in KO mice compared to WT controls (Figure 4, I-L). Together, these findings demonstrate that HDAC11 deficiency has a general salutary impact on mouse metabolic health in the context of acute and chronic fat and caloric excess.

**Figure 4.**
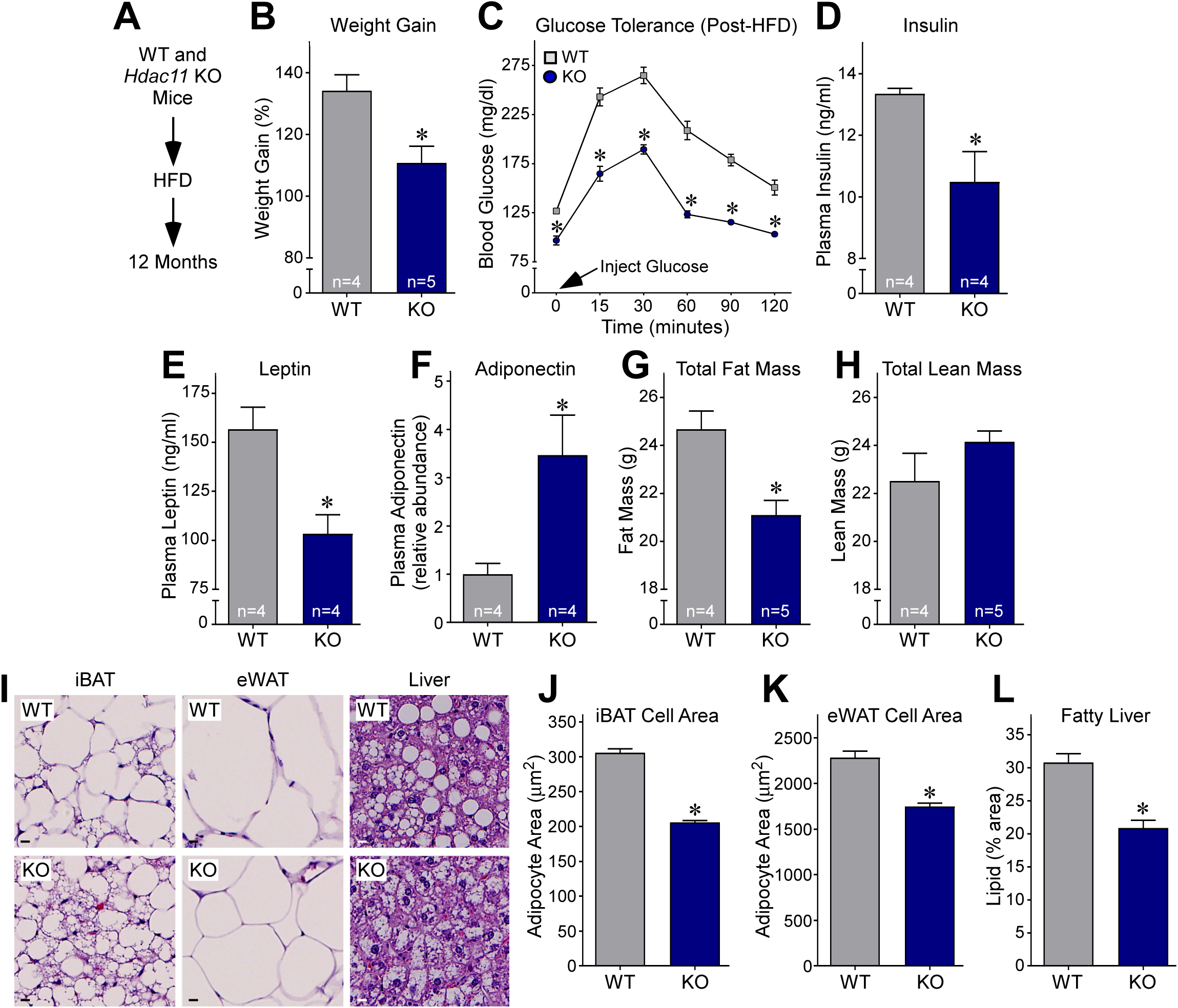
HDAC11 deficiency protects against deleterious effects of chronic high fat feeding. (**A**) Schematic representation of the experiment. (**B**) Weight gain (%) was calculated at 12 months. (**C**) Glucose tolerance was assessed in mice following 12 months of high fat feeding; n = 4-5/group. (**D-E**) Plasma insulin and leptin concentrations were determined by ELISA. (**F**) Plasma adiponectin concentrations were determined via immunoblotting and normalized to IgG levels. (**G-H**) Body composition (total fat and lean mass) was determined using QMRI. (**I**) H&E staining of iBAT, eWAT and liver following 12 months of high fat feeding. Scale bar:10 μm. (**J-L**) Adipocyte area in iBAT and eWAT, and lipid accumulation in liver was quantified as described in the Methods section. Values for all graphs represent means +SEM; \**P*<0.05 vs. WT. Two-way ANOVA with Sidak’s multiple comparisons test for four groups was employed for (**C**); Student’s t-test was used for all other graphs.

### HDAC11 knockdown promotes brown adipocyte differentiation

A loss-of-function approach was employed to address the possibility that HDAC11 functions in a cell autonomous manner to suppress brown adipocyte differentiation. Initially, shRNA was employed to diminish expression of endogenous HDAC11 in mouse embryonic fibroblasts (MEFs), and the cells were subsequently exposed to BAT differentiation medium for seven days (Figure 5A). Reduced HDAC11 expression correlated with enhanced differentiation of MEFs into adipocyte-like cells harboring multilocular lipid droplets that are characteristic of BAT (Figure 5, B-D). Furthermore, HDAC11 knockdown led to a dramatic enhancement of *Ucp1* mRNA and protein expression, which correlated with augmented *Pgc1α* mRNA and protein levels (Figure 5, E-G).

**Figure 5.**
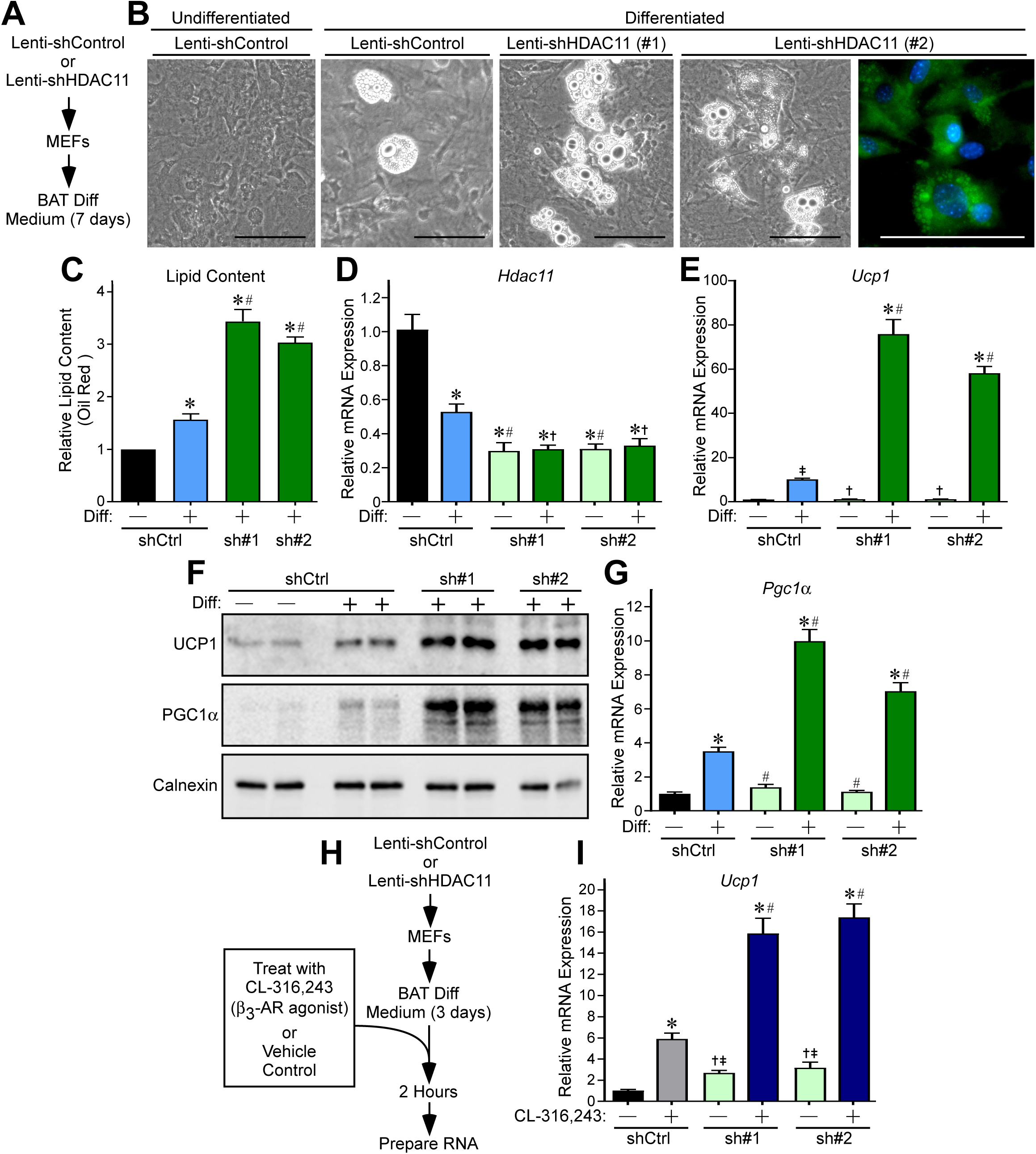
HDAC11 knockdown enhances differentiation of MEFs into brown adipocyte-like cells. (**A**) Schematic representation of the MEF differentiation assay. (**B**) Phase contrast images of MEFs differentiated into brown adipocyte-like cells. BODIPY staining shows multilocular lipid droplets characteristic of brown adipocytes. Scale bar: 100 μm. (**C**) Lipid content was quantified using colorimetric Oil Red O staining; n = 4 plates/condition. (**D-E**) Quantitative PCR analysis of *Hdac11 and Ucp1* mRNA expression; n = 4 plates of cells/condition. (**F**) Immunoblot analysis of UCP1 and PGC1α protein expression; each lane represents protein from an independent plate of cells. Calnexin served as a loading control. (**G**) Quantitative PCR analysis of *Pgc1*α mRNA expression; n = 4 plates of cells/condition. (**H**) Schematic representation of the β_3_-AR signaling assay. (**I**) Quantitative PCR analysis of *Ucp1* mRNA expression following 2 hours of treatment of cells with CL-316,243 or vehicle control; n = 4 plates of cells/condition. Values for all graphs represent means +SEM; \**P*<0.05 vs. shControl undifferentiated or unstimulated; ^#^*P*<0.05 vs. shControl differentiated or stimulated using ANOVA with Tukey’s multiple comparisons test. ^†^*P*<0.05 vs shControl differentiated or stimulated; ^‡^*P*<0.05 vs shControl undifferentiated or unstimulated using Student’s t-test following significant ANOVA.

Next, an acute study was performed to address whether HDAC11 regulates intrinsic responsiveness of cells to β_3_-AR signaling (Figure 5H). Consistent with the findings from the *ex vivo* study of BAT (Figure 2B), β_3_-AR agonist-mediated *Ucp1* induction was substantially increased in partially differentiated MEFs in which HDAC11 expression was attenuated using shRNA (Figure 5I).

### Repression of brown adipocyte differentiation by HDAC11 is dependent on association with BRD2

A prior proteomics study suggested that HDAC11 is capable of associating with BRD2, a member of the BET family of acetyl-histone binding proteins that has been implicated as a negative regulator of brown adipocyte differentiation (34;35). Thus, we hypothesized that HDAC11-mediated control of BAT differentiation is dependent on its interaction with BRD2. To initially address this hypothesis, experiments were performed to confirm the association between HDAC11 and BRD2. Confocal microscopy of HIB1B pre-brown adipocytes ectopically expressing tagged forms of HDAC11 and BRD2 revealed that, when expressed individually, the proteins were pan-cellular and nuclear localized, respectively (Figure 6, A and B). However, co-expression of the proteins led to re-distribution of HDAC11 exclusively to the nuclear compartment, where it co-localized with BRD2 (Figure 6, C-F).

**Figure 6.**
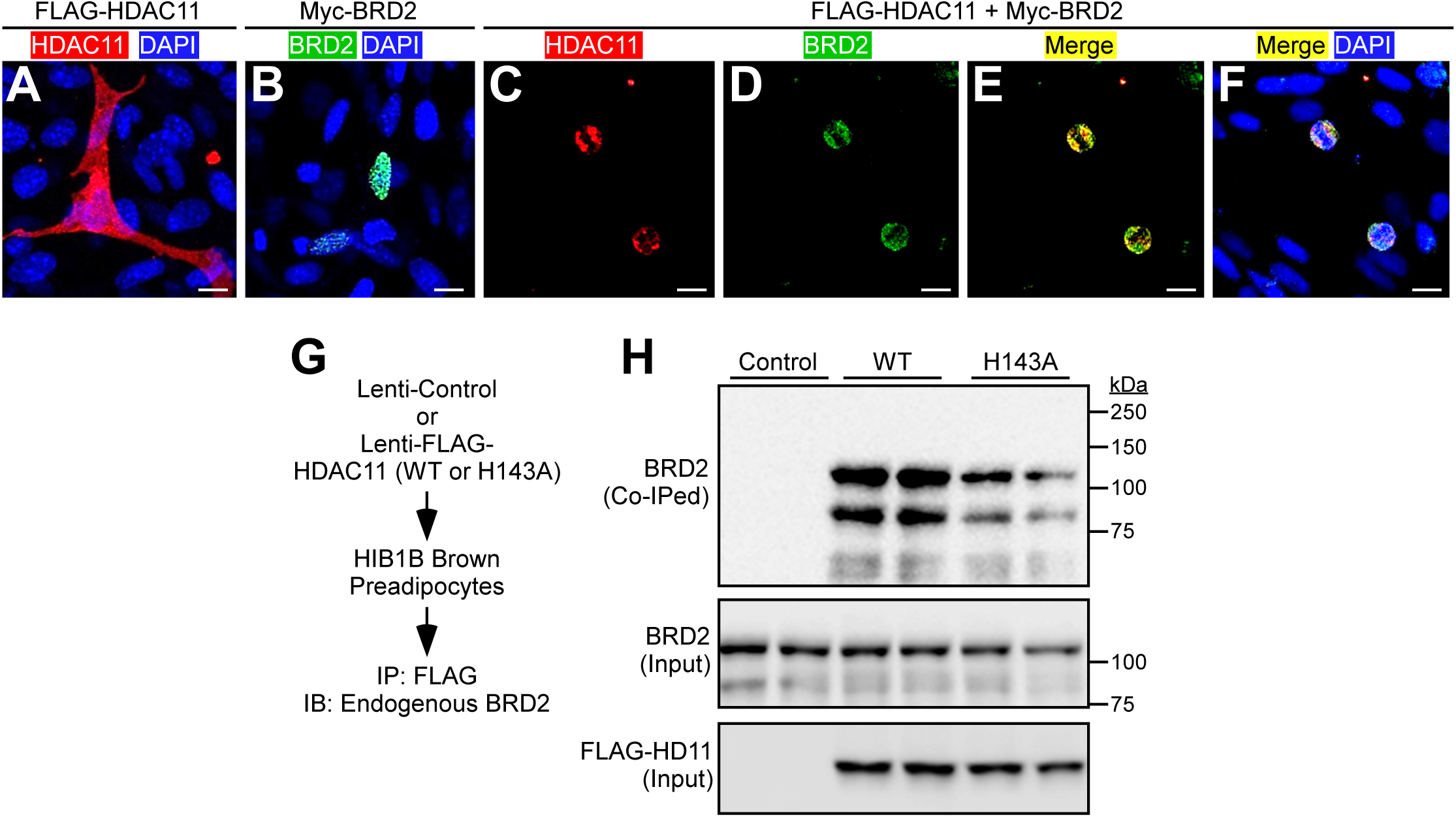
HDAC11 associates with BRD2. (**A-F**) Confocal imaging of epitope-tagged HDAC11 and BRD2 in transiently transfected HIB1B preadipocytes. (**G**) Schematic representation of the HDAC11:BRD2 co-immunoprecipitation (IP) assay. (**H**) Immunoblot (IB) assessment of endogenous BRD2 co-immunoprecipitating with ectopic HDAC11. Total amounts of HDAC11 and BRD2 in protein homogenates were also determined (input).

To further validate that HDAC11 and BRD2 associate within a common regulatory complex, co-immunoprecipitation (Co-IP) assays were performed with cells ectopically expressing wildtype HDAC11 (WT) or a catalytically inactive version of HDAC11 harboring a histidine-to-alanine substitution at position 143 (H143A) within the deacetylase domain (Figure 6G). Endogenous BRD2 was effectively co-IPed with HDAC11 (both WT and H143A) in HIB1B cells and HEK293 cells (Figure 6H and data not shown).

Employing a series of deletion constructs in transiently transfected HEK293 cells, the region between amino acids 200 and 250 within the catalytic domain of HDAC11 was determined to be critical for association with BRD2, while the extraterminal (ET) domain of BRD2 was found to be required for association with HDAC11 (Figure 7, A-F). Of note, the related BET family member, BRD4, failed to associate with HDAC11, which suggests that HDAC11 selectively interacts with BRD2 as opposed to generally targeting BET proteins (Supplemental Figure 2).

**Figure 7.**
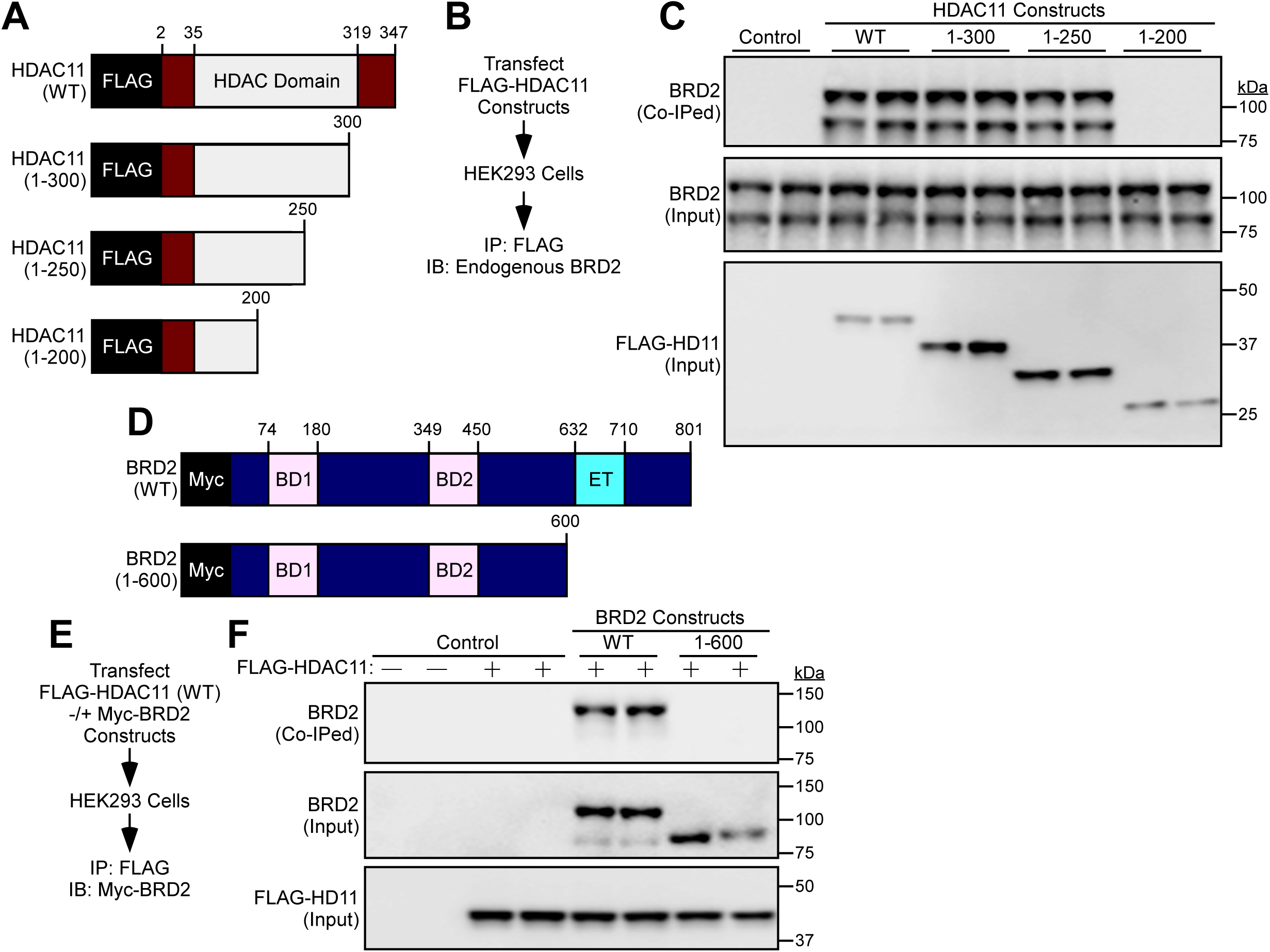
Mapping of HDAC11:BRD2 association domains. (**A**) Schematic representation of human HDAC11 and carboxy-terminal deletion constructs. (**B**) Schematic representation of the HDAC11:BRD2 co-immunopreciptation (IP) assay with HDAC11 deletion constructs. (**C**) Immunoblot (IB) assessment of endogenous BRD2 co-immunoprecipitating with ectopic HDAC11. Total amounts of HDAC11 and BRD2 in protein homogenates were also determined (input). (**D**) Schematic representation of human BRD2 and a carboxy-terminal deletion construct. BD1, bromodomain 1; BD2, bromodomain 2; ET, extra-terminal domain. (**E**) Schematic representation of the HDAC11:BRD2 co-immunoprecipitation assay with ectopic HDAC11 and ectopic BRD2. (**F**) Immunoblot assessment of ectopic BRD2 co-immunoprecipitating with ectopic HDAC11. Total amounts of epitope-tagged HDAC11 and BRD2 in protein homogenates were also determined (input).

Having confirmed the association between HDAC11 and BRD2, we next sought to determine whether BRD2 contributes to HDAC11-mediated regulation of brown adipocyte differentiation. Wild-type HDAC11 was ectopically overexpressed in HIB1B brown preadipocytes in the absence or presence of shRNA targeting BRD2, and the cells were subsequently exposed to BAT differentiation medium for 4 days (Figure 8A). Overexpression of HDAC11 WT significantly reduced *Ucp1* and *Pgc1α* mRNA expression upon HIB1B differentiation (Figure 8, B and C). In contrast, catalytically inactive HDAC11 failed to repress expression of *Ucp1* or *Pgc1α* (Supplemental Figure 3). Remarkably, BRD2 knockdown completely blocked the ability of HDAC11 to suppress expression of *Ucp1* and *Pgc1α* mRNA and protein expression, and also rescued expression of PPAR_γ_ and C/EBPα (Figure 8, B-E). These findings suggest that HDAC11 inhibits BAT differentiation and thermogenic gene expression, at least in part, through association with BRD2 (Figure 8F).

**Figure 8.**
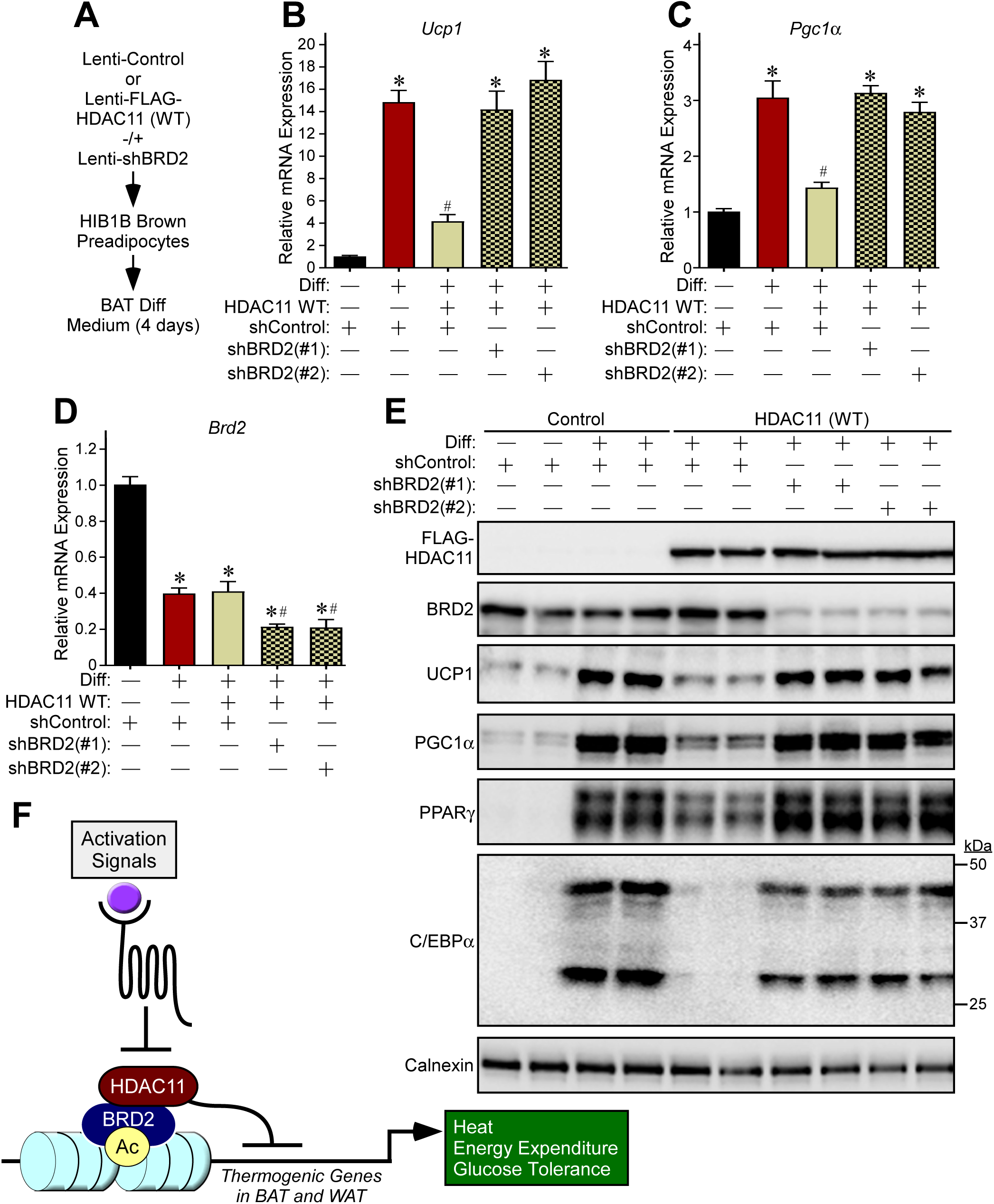
HDAC11-dependent suppression of brown adipocyte differentiation requires BRD2. (**A**) Schematic representation of the experiment. (**B-D**) Quantitative PCR analysis of *Ucp1*, *Pgc1*α and *Brd2* mRNA expression; n = 4 plates of cells/condition. (**E**) Immunoblot analysis of the indicated proteins; each lane represents protein from an independent plate of cells. Calnexin served as a loading control. Values for all graphs represent means +SEM; ^*^*P*<0.05 vs. shControl undifferentiated; ^#^*P*<0.05 vs. shControl differentiated. Two-way ANOVA with Tukey’s multiple comparisons test was employed. (**F**) A model for HDAC11-mediated control of thermoregulation and energy homeostasis.

## Discussion

Here, we describe a role for an obscure lysine deacetylase, HDAC11, in the transcriptional control of thermogenic gene expression in adipose tissue through association with BRD2, an acetyl-lysine binding epigenetic ‘reader’ protein. HDAC11 deficient mice are healthy and phenotypically indistinguishable from WT controls, yet are resistant to cold challenge and the adverse metabolic consequences of high fat feeding due to enhanced metabolic activity in BAT, and increased beiging of WAT. The findings reveal a highly druggable transcriptional pathway for the regulation of energy expenditure, and thus suggest novel approaches for combatting the world-wide pandemics of obesity and T2D via selective inhibition of HDAC11, or disruption of HDAC11:BRD2 association.

Increased thermogenesis and energy expenditure in HDAC11-deficient mice is associated with augmented BAT formation and function, and beiging of WAT. Although we cannot rule out the possibility that HDAC11 controls adipose tissue through indirect mechanisms, such as by enhancing sympathetic drive, evidence is presented to suggest that HDAC11 functions in a cell autonomous manner to control thermogenic gene expression in adipose tissue. For example, BAT explants from KO mice exhibit dramatically enhanced responsiveness to acute CL-316,243 stimulation compared to WT controls, despite equivalent expression of the β_3_-AR (Figure 2). Furthermore, knockdown of HDAC11 expression in fibroblasts is sufficient to promote *Ucp1* and *Pgc1α* expression and amplify β_3_-AR-stimulated thermogenic gene expression (Figure 5), and ectopic expression of HDAC11 in preadipocytes has the converse actions (Figure 8). Future studies with mice in which HDAC11 is conditionally deleted in a tissue-specific manner will elucidate the relative contributions of HDAC11 in distinct cell types to the control of systemic metabolism.

Suppression of thermogenic gene expression by HDAC11 is due, at least in part, to its association with BRD2 (Figure 8). However, HDAC11 deletion does not completely phenocopy BRD2 hypomorphic mice engineered using gene trap mutagenesis (35). In these animals, diminishing global BRD2 expression by ~50% resulted in increased BAT formation and thermogenesis and reduced blood glucose in the fasting and fed states, consistent with what was observed in HDAC11 KO mice. However, disruption of BRD2 expression in mice also led to extreme obesity, which contrasts with the lean phenotype of HDAC11-deficient animals. These data suggest that HDAC11 selectively regulates only a subset of BRD2 target genes, and that inhibition of HDAC11 catalytic activity or interference of HDAC11 association with BRD2 would be better tolerated than general inhibition of BRD2 with compounds such as bromodomain inhibitors. Given that HDAC11 can also localize to the cytoplasmic compartment (Figure 6A), and has been shown to associate with a multitude of extra-nuclear proteins, including splicing factors (36), it remains possible the HDAC11 also governs thermogenic gene expression through BRD2-independent and non-genomic actions.

The 18 mammalian HDACs are clustered into four classes: class I HDACs (1, 2, 3 and 8), class II HDACs (4, 5, 6, 7, 9 and 10), class III HDACs (SirT 1 - 7) and class IV (HDAC11) (33). Class I, II and IV HDACs are zinc-dependent enzymes, while class III HDACs, which are also known as sirtuins, require NAD^+^ as a co-factor for catalytic activity. Sirtuins are most commonly associated with regulation of metabolism (37), although zinc-dependent HDACs have recently emerged as regulators of thermogenic gene expression. For example, HDAC1 has been shown to associate with regulatory elements of BAT-specific genes, resulting in reduced histone H3K27 acetylation and consequent transcriptional repression (38). Another class I HDAC, HDAC3, has also been linked to the regulation of BAT gene expression and thermogenesis in mice (39-41). The class II HDAC, HDAC9, has been demonstrated to repress adipogenesis, and deletion of HDAC9 in mice is sufficient to promote beiging and increase energy expenditure and adaptive thermogenesis in the context of high fat feeding (42-44). However, unlike HDAC1 and HDAC3, HDAC9-mediated regulation of adipogenic gene expression is not dependent on its deacetylase domain, and instead is mediated by the amino-terminal co-factor interaction module of the protein (42).

HDAC11 enzymatic activity appears to be required for inhibition of adipocyte gene expression, as evidenced by the failure of catalytically inactive HDAC11 (H143A) to effectively suppress expression of *Ucp1* and *Pgc1α* in HIB1B cells (Supplemental Figure 3). This finding supports the concept of employing selective small molecule inhibitors of HDAC11 to increase energy expenditure, and the feasibility of this therapeutic approach is bolstered by the fact that four HDAC inhibitors are FDA approved drugs (45). Nonetheless, pan-HDAC inhibition is accompanied by untoward toxicities, such as thrombocytopenia, nausea and fatigue (46). We posit that selective inhibition of HDAC11 would provide a safer and more effective alternative to pan-HDAC blockade. This notion is supported by the observation that global deletion of HDAC11 in mice is extremely well tolerated, while whole body removal of the previously mentioned HDACs that are implicated in the control of obesity and metabolism results in embryonic lethality or heart failure (47-50).

A recent discovery highlights the attainability of selective HDAC11 inhibitors. Unlike other HDACs, HDAC11 is a weak histone deacetylase, and instead is a highly effective lysine defatty-acylase; HDAC11 defatty-acylates substrates with an efficiency that is >10,000-fold greater than its deacetylase activity (51-53). This finding suggests that HDAC11 functions, at least in part, in a manner analogous to sirtuins, which have the ability to proficiently catalyze removal of acyl groups in addition to acetyl, as opposed to other zinc-dependent HDACs, which primarily target acetyl moieties on lysine (54). The combination of this unique catalytic activity among the zinc-dependent HDACs and the divergent phylogeny of HDAC11 underscore the potential to selectively target HDAC11 with small molecule inhibitors.

An alternative approach to alleviate HDAC11-mediated suppression of thermogenesis would be to inhibit its interaction with BRD2. In this regard, a region of BRD2 that encompasses the ET domain was found to be required for association with HDAC11 (Figure 7F). ET domains of BET family members control gene transcription by functioning as interaction modules for co-factors such as the NSD3 histone methyltransferase (55). Recent NMR-derived structural analyses revealed compact motifs governing ET domain binding with partner proteins, suggesting the feasibility of employing small molecule ligands to disrupt ET domain-dependent functions to achieve therapeutic benefit (56;57).

In conclusion, the present work uncovered novel insights into the molecular underpinnings on thermogenic gene regulation and the control of metabolism via BAT and beiging of WAT. HDAC11-deficient mice are healthy and exhibit dramatic amelioration of multiple metabolic parameters in mouse models of obesity. Given the unique features of HDAC11 highlighted above, the current findings provide unprecedented avenues for the development of innovative epigenetic therapies for obesity and obesity-related metabolic disorders by targeting the HDAC11:BRD2 axis.

## Methods

### Animals

Constitutive HDAC11KO allele mice on a C57BL/6J background were provided by Merck Research Laboratories and generated by a targeted deletion of floxed exon 3 of the HDAC11 gene utilizing Rosa26 promoter-driven cre-recombinase expression. The Cre transgene was removed by segregation. These mice are now available from Taconic (Model 6978). HDAC11 heterozygous (+/-) mice were bred to obtain HDAC11 KO mice (-/-) and their WT (+/+) littermate controls. Genotypes were confirmed via PCR using specific primers (Supplemental Table 3). Mice were maintained in a temperature (22^0^C) in a light-controlled vivarium with free access to water and standard rodent chow. All animal studies were initiated with 10-week-old male mice.

### Human tissue biopsies

Brown adipose tissues were collected from deep neck regions of patients undergoing thyroid surgeries at Florida Hospital. Subcutaneous abdominal white adipose tissue samples were obtained from participants of a cross-sectional study conducted at the Florida Hospital Translational Research Institute for Metabolism and Diabetes. Tissues were immediately snap-frozen and stored in liquid nitrogen. Tissue samples used in this study are a subset of those described in a prior report (Supplemental Table 4) (58).

### Cold challenge and high fat feeding

For cold exposure experiment, WT and HDAC11 KO mice were kept at 4°C for 24h in indirect calorimetry metabolic cages (CLAMS; Columbus Oxymax metabolic monitoring system). Core body temperature was monitored every two hours for the duration of the study using a rectal probe (Physitemp Instruments Inc.) connected to a physiological monitoring unit (THM150; VisualSonics). Mice were fed a high fat diet (HFD; Research Diets Inc., D09071702, 58% calories derived from fat) for 8 days or 12 months. Metabolic monitoring of mice in response to acute HFD was performed using CLAMS. For glucose tolerance tests, mice were fasted for 6h prior to administration of glucose (2mg/g body weight) via intraperitoneal injection. Tail vein blood was collected, and blood glucose measured using a commercial glucometer (Bayer Contour) at indicated time points. Body composition was determined using MRI (EchoMRI, Houston, TX, USA) to assess fat and lean mass. Mice were sacrificed isoflurane inhalation followed by cervical dislocation, blood was collected via cardiocentesis, and adipose tissues and liver were harvested following tissue perfusion with saline.

### Histological analyses

Adipose and liver tissues were formalin fixed and embedded in paraffin. 5μm-thick tissue sections were deparaffinized, hydrated and processed for hematoxylin and eosin staining using conventional methods. Sections were then dehydrated and mounted using Organo/Limonene Mount (Sigma-Aldrich). 5μm-thick tissue sections of ingWAT from HDAC11 KO and WT mice exposed to cold challenge were deparaffinized, hydrated and subjected to antigen retrieval using citrate buffer (Vector Laboratories). Endogenous peroxidase activity was inhibited using BLOXALL blocking solution (Vector Laboratories) for 10 minutes, and sections were incubated in 5% BSA for 1hr at RT. Tissue sections were incubated overnight with anti-UCP1 antibody (Abcam) followed by incubation with biotinylated secondary antibody. Immunoperoxidase detection was performed using commercially available reagents as per the manufacturer’s instructions (Vector Laboratories). Images were acquired using an Olympus microscope (BX51) equipped with a multicolor camera (DP72). To quantify adipocyte size, 5 μm thick sections per paraffin embedded tissue block were stained with H&E and imaged at 20x magnification. High-resolution images (4-5 fields per section per animal) were acquired for adipocyte morphometry analysis. Automated analysis of adipose tissue cellularity was performed using Adiposoft open-source software (59). Average adipocyte areas were calculated for each animal, grouped and plotted as mean adipocyte area for each genotype.

### Quantification of circulating insulin, leptin and adiponectin levels

Blood was collected in 1.5ml microfuge tubes containing EDTA and stored on ice. Tubes were centrifuged at 14000rpm for 5 minutes at 4°C, and supernatants were transferred to new microfuge tubes and stored at - 80°C. Samples were used to determine levels of plasma insulin and leptin by ELISA according to the manufacturer’s instructions (CrystalChem). Plasma adiponectin estimation was performed by immunoblotting with adiponectin-specific antibody (Acrp30) followed by quantification of adiponectin-positive bands by densitometry, as previously described (60).

### Ex vivo BAT assay

iBAT was collected from 10-week old male mice. Each BAT explant was dissected in half and placed in medium (5 mls of DMEM supplemental with 10% FBS) in sterile 14 ml round bottom polystyrene tubes. Immediately thereafter, the explants were treated with CL-316243 (10 μM, Cayman) or vehicle control (DMSO, 0.1% final concentration), and incubated in at 37°C for 3 hours in a cell culture incubator. The tissues were flash frozen in liquid nitrogen prior to RNA extraction.

### qRT-PCR

RNA was prepared from cells and adipose tissues using QIAzol lysis reagent (Qiagen). 500ng of total RNA was reverse-transcribed and cDNA synthesized using Verso cDNA synthesis kit (Thermo Fisher Scientific). qPCR was performed on a StepOnePlus Real-Time PCR System (Applied Biosystems) using PowerUp™ SYBR Green Master Mix (Thermo Fisher Scientific) and specific primers (Supplemental Table 1). Amplicon abundance was calculated using the 2^−ΔΔCT^ method, and normalized to 18S RNA or *Hprt1*. For analysis of mRNA expression in human tissue biopsies, RNA was extracted using commercial reagents (RNeasy Lipid Tissue Kit; Qiagen) and qPCR was performed using Taqman Fast Virus 1-step Master reaction mix (Life Technologies) on a ViiAR 7 Real-Time PCR system (Applied Biosystems). Pre-designed primer probe sets were used (Hs00978041_m1 for *Hdac11*, Hs02800695_m1 for *Hprt1*; Life Technologies). Relative gene expression data were calculated using the ΔC_t_.

### Immunoblotting

Total protein was isolated from undifferentiated or brown adipocyte-differentiated cells (HIB1B or MEFs) using lysis buffer comprised of NaCl (300 mM) and TritonX-100 (0.5% in PBS pH 7.4) supplemented with protease and phosphatase inhibitor cocktail (Thermo Fisher Scientific). Proteins were separated using SDS-PAGE and transferred onto nitrocellulose membranes (0.45μm; Life Science Products). Blots were probed with primary antibodies specific for UCP1 (Abcam), PGC-1α (Calbiochem), C/EBPα, PPARγ, BRD2 (Cell Signaling), FLAG M2-HRP (Sigma), calnexin or α-tubulin (Santa Cruz Biotechnology) overnight at 4°C. Following incubation with primary antibodies, blots were then probed with appropriate HRP-conjugated mouse or rabbit secondary antibodies (Southern Biotech). Protein bands were visualized using enhanced chemiluminescent HRP substrate (SuperSignal West Pico Chemiluminescent Substrate; Thermo Scientific) on a FluorChem HD2 Imager (Alpha Innotech).

### Plasmids

pcDNA3.1(+)-based plasmids encoding human HDAC11, human BRD2 and deletion constructs fused in-frame to an amino-terminal epitope tags were generated using PCR with PFU Turbo polymerase (Agilent Technologies) and specific primers (Supplemental Table 2). Site-directed mutagenesis to create pcDNA3.1 encoding human HDAC11 (H143A) was performed using the QuickChange method (Agilent Technologies). For lentivirus production, complementary DNAs encoding HDAC11 were cloned into pLenti CMV Hygro DEST. pLKO.1 plasmids (Sigma) encoding shRNA for murine HDAC11 (TRCN0000339501 and TRCN0000377043), BRD2 (TRCN0000023960 and TRCN0000023962) and a negative control (SHC002) were obtained through Functional Genomics Facility at the University of Colorado Cancer Center.

### Lentivirus production

Lentiviruses were generated by co-transfecting L293 cells with pLenti or pLKO.1 vectors in combination with packaging plasmid (psPAX2) and envelope plasmid (pMD2.G). Virus-containing cell culture supernatants were collected, filtered and used for experiments. pLenti-CMV-Hygro DEST (w117-1) was a gift from Eric Campeau & Paul Kaufman (Addgene). psPAX2 and pMD2.G were gifts from Didier Trono (Addgene). Virus-containing cell culture supernatants were collected, filtered (0.45 μm; Life Science Products) and frozen at −80°C until needed.

### Cell culture and differentiation

Mouse embryonic fibroblasts (MEFs) were isolated from C57BL/6 E14.5 embryos using enzymatic digestion as described and maintained in DMEM/Hi glucose (Hyclone) supplemented with 10% fetal bovine serum (FBS; Gemini), 1% Penicillin– Streptomycin (Gibco) and 1.1% GlutaMAX (Gibco) (61). For differentiation into brown adipocytes, 4×10^5^ cells were seeded per 60-mm dish and cultured in DMEM medium containing dexamethasone (1 μM), isobutylmethylxanthine (0.5 mM), T_3_ (50 nM), insulin (5 μg/ml), and rosiglitazone (2.5 μM) for 48 hours after lentivirus-mediated knockdown of HDAC11 (shHDAC11, or control shRNA). Differentiated cells were then maintained in media supplemented with insulin (5 μg/ml) and rosiglitazone (0.5 μM). Cells were treated with a β_3-_AR agonist, CL-316,243 (1 μM; Cayman), for 3 hours. Differentiated MEF cultures were stained with 0.5% Oil Red O to determine lipid content. Lipid droplets were visualized using BODIPY™493/503 (2μM; 4,4-Difluoro-1,3,5,7,8-Pentamethyl-4-Bora-3a,4a-Diaza-s-Indacene; Molecular Probes). HIB1B brown preadipocytes were maintained in DMEM with 10% FBS and differentiated into adipocytes by the addition of rosiglitazone (1 μM) for 4 days (62). For experiments with lentiviruses, 3mL of filtered viral supernatant supplemented with 3 μL polybrene (Sigma) was added to cells, and the medium was changed 24 hours post-infection. For MEFs, cells were grown to confluency (48 hours), the medium was replenished, and cells were maintained in a growth arrested state for an additional 48 hours prior to induction of differentiation. For HIB1B adipocytes, cells were infected at ~60% confluency (24 hours), the medium was subsequently replenished, and cells recovered for 24 hours prior induction of differentiation.

### Co-Immunoprecipitation

HIB1B brown preadipocytes were infected with lentiviruses encoding FLAG-tagged wildtype human HDAC11 or catalytically inactive mutant form (H143A). HEK293 cells were transfected with tagged cDNA expression vectors encoding human HDAC11 (FLAG-tagged), deletion constructs of HDAC11 (FLAG-tagged), human wildtype BRD2 (Myc-tagged) or ET domain deleted BRD2 (Myc-tagged) using polyethylenimine (PEI) for 48 hours. Cells were lysed in buffer containing Tris (50 mM, pH 7.5), 150 mM NaCl and Triton X-100 (0.5 %) using a syringe with a 25-gauge needle. Total protein was collected after centrifugation of lysates at 13,000 rpm at 4°C for 20 min. Protein homogenates (0.5 mg) were diluted in equilibration buffer (50 mM Tris, 150 mM NaCl, pH 7.4; 500 μl total volume) and were immunoprecipitated using anti-FLAG IP resin (25 μl packed resin, GenScript), overnight at 4°C on a rotator, washed (three times with equilibration buffer; 0.5 mL per wash), denatured in sample loading buffer and resolved through 10% SDS polyacrylamide gels. Proteins were transferred to nitrocellulose membranes (0.45 μm; Life Science Products) and immunoblotted as described above using anti-BRD2 (Cell Signaling Technologies), anti-Myc (Santa Cruz Biotechnology) or anti-BRD4 (Bethyl Laboratories) antibodies. Whole-cell lysates were used as input controls.

### Indirect immunofluorescence

HIB1B brown preadipocytes grown on coverslips were transiently transfected with pcDNA3.1-based plasmids encoding human HDAC11 (FLAG-tagged) and human BRD2 (Myc-tagged) for 48 hours using Lipofectamine 3000 (Life Technologies). Cells were fixed with 4% paraformaldehyde at room temperature for ten minutes, followed by permeabilization using PBS-T (0.2% Triton X-100) for fifteen minutes with gentle rocking. Cells on coverslips were blocked with PBS containing 5% BSA for 30 minutes at room temperature and incubated with primary antibodies (FLAG, Sigma, 1:200; c-Myc, Cell Signaling, 1:200) for 2 hours. Secondary antibodies (Anti Rabbit Alexa Fluor® 488, 1:800; Anti Mouse Alexa Fluor® 555, 1:800; Molecular Probes) were applied for 30 minutes at room temperature. Coverslips were mounted on glass slides using anti-fade mounting reagent with DAPI (Vector Laboratories, H-1200). Images were obtained on an Olympus FLUOVIEW FV1000 confocal laser scanning microscope (UC Denver, Advanced Light Microscopy Core) and processed through FV-Viewer software (Olympus).

### Statistics

Statistical significance (*P*<0.05) was determined using unpaired t-test (two groups) or two-way ANOVA with correction for multiple comparisons via a Tukey post hoc test (GraphPad Prism 7.02).

### Study approval

Animal studies were conducted using a protocol approved by the Institutional Animal Care and Use Committee of the University of Colorado Anschutz Medical Campus, following appropriate guidelines. Studies involving human subjects were approved by the Florida Hospital Institutional Review Board, and participants gave written informed consent.

## Author contributions

R.A.B., B.S.F., M.S.S., T.H., M.A.C., Y.H.L., and M.F.P. performed the research and analyzed data. L.S., D.L., P.L., K.S., P.E.S., S.C., and E.S. provided reagents, designed experiments and analyzed data. L.M.S., S.R.S., and T.A.M. designed and supervised the research and analyzed data. R.A.B. and T.A.M. wrote the manuscript.

## Acknowledgements

We thank Matthew Jackman for assistance with metabolic phenotyping studies at the Colorado Nutrition & Obesity Research Center (NORC) Animal Satellite Facility, Kelley Brodsky for assistance with microscopy, and Sara Wennersten, Katharina Lutter and Korey Haefner for help with *in vivo* studies. We are grateful to Dwight Klemm and Keith Koch for critical input. This work was supported by the National Institutes of Health (HL116848, HL127240 and AG043822) and the American Heart Association (16SFRN31400013) to T.A.M. R.A.B. received funding from the Canadian Institutes of Health Research (FRN-216927). M.S.S. was funded by a T32 training grant and an F32 fellowship from the National Institutes of Health (T32HL007822 and F32HL126354). Y.H.L. was supported by an American Heart Association postdoctoral fellowship (16POST30960017).

## Supplemental information

Supplemental Information includes three figures and three tables and can be found with this article online.

